# Entrainment echoes in the cerebellum

**DOI:** 10.1101/2024.03.06.583255

**Authors:** Benedikt Zoefel, Omid Abbasi, Joachim Gross, Sonja A Kotz

**Affiliations:** Centre de Recherche Cerveau et Cognition (CerCo), CNRS, Toulouse, France; Université Paul Sabatier Toulouse III, France; Institute for Biomagnetism and Biosignal Analysis, University of Münster, Germany; Otto-Creutzfeldt-Center for Cognitive and Behavioral Neuroscience, University of Münster, Germany; Department of Neuropsychology & Psychopharmacology, Faculty of Psychology and Neuroscience, Maastricht University, the Netherlands; Department of Neuropsychology, Max Planck Institute for Human Cognitive and Brain Sciences, Leipzig, Germany

## Abstract

Historically, researchers have considered the cerebellum a coordinator of motor programs that ensures precise timing of movements and their adaptation to external events^1^. However, it has become increasingly clear that this role is not restricted to the motor system^2^. Rather, the cerebellum seems to play an important role in temporal prediction in general, as shown in its involvement in multiple functions that rely on precise event timing^3^. Although previous work suggested that the cerebellum exclusively predicts the interval between two events^4^, rather than tracking a global rhythm, it is also active when a rhythmic stimulus *changes* in rate^5^. The latter finding is in line with a cerebellar role in speech processing^6^ that entails frequent rate changes^7^. Neural mechanisms underlying the cerebellum’s involvement in speech processing, however, remain poorly understood. Moreover, there is a lack of studies contrasting speech and non-speech stimuli to establish speech-specificity of the observed effects^8^. In a re-analysis of magnetoencephalography (MEG) data^9^, we found that activity in the cerebellum aligned to rhythmic sequences of noise-vocoded speech, irrespective of its intelligibility. We then tested whether these “entrained” responses persist, and how they interact with other brain regions, when the rhythmic stimulus stopped and temporal predictions had to be updated. We found that only intelligible speech produced rhythmic responses in the cerebellum that outlasted the stimulus. During this “entrainment echo”, but not during rhythmic speech itself, cerebellar activity was coupled with that in the left inferior frontal gyrus (IFG), and specifically at rates corresponding to the preceding stimulus rhythm. This finding represents unprecedented evidence for specific cerebellum-driven temporal predictions in speech processing and their relay to cortical regions.

## Results

### Overview

Participants listened to rhythmic sequences of noise-vocoded, monosyllabic words (Fig. 1A) that were presented at 2 Hz or 3 Hz and either intelligible (16-channel speech) or unintelligible and noise-like (1-channel speech). The main aim of the original work^9^ was to test for the existence of “entrainment echoes”, rhythmic brain responses that are produced by a rhythmic stimulus and persist for some cycles after its offset^9–11^. Endogenous neural oscillations play a critical role in the field of “neural entrainment”^12^, but are difficult to demonstrate *during* a rhythmic stimulus, as multiple alternatives can explain a rhythmic neural response^13^. Entrainment echoes are measured immediately *after* the rhythmic stimulus and therefore support the involvement of endogenous oscillations, which should, once entrained, outlast the stimulus before going back to their natural state^14,15^. As the offset of the rhythmic stimulus violates temporal expectation (induced by the stimulus rhythm), entrainment echoes might also give insight into neural mechanisms underlying participants’ prediction of upcoming events^10^.

**Figure 1.**
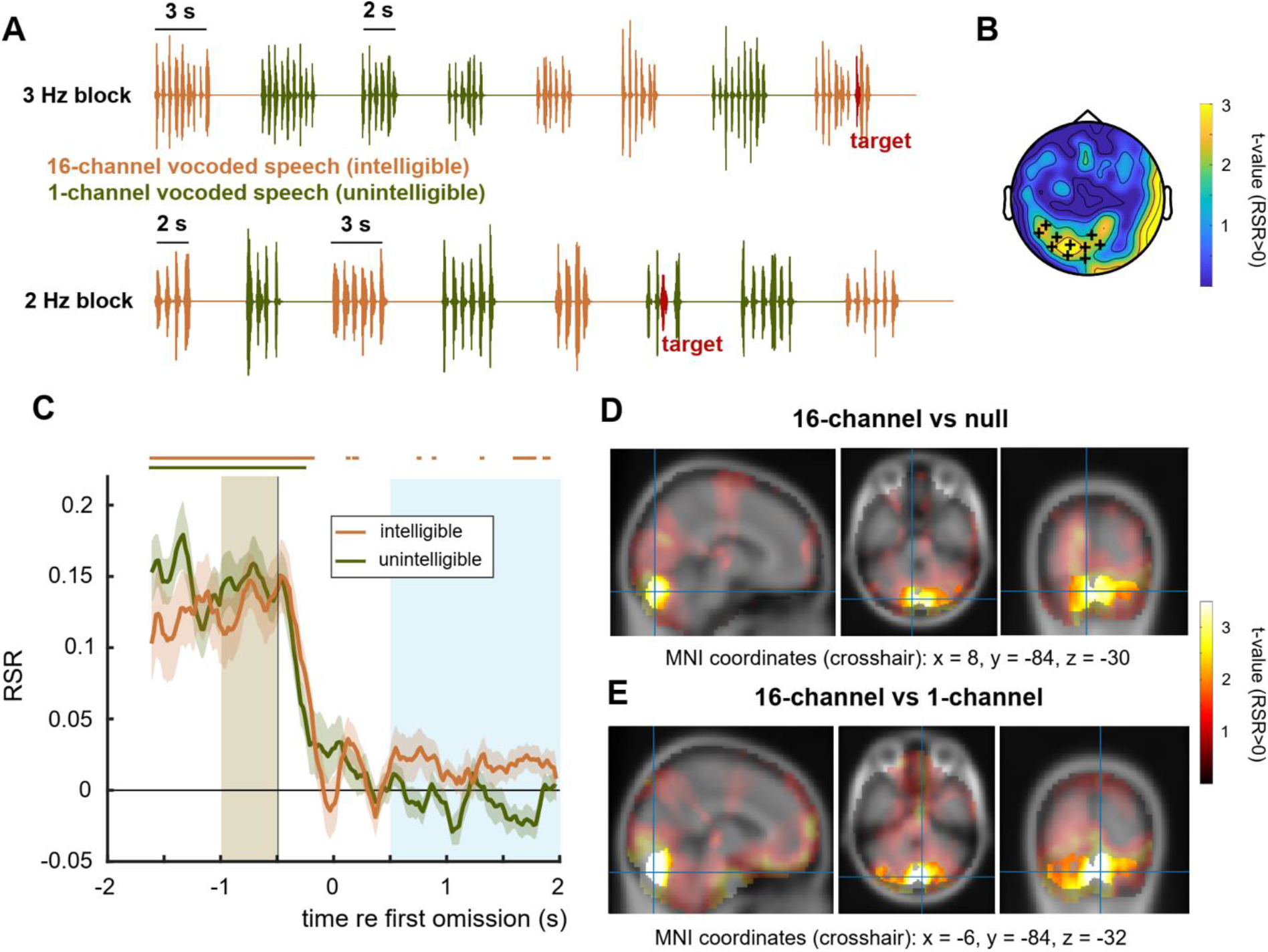
Paradigm, previous results, and source-localised entrainment echo. **A.** Experimental paradigm. Participants listened to rhythmic sequences of intelligible or unintelligible speech and were asked to detect rhythmic irregularities (red). **B.** The original study^9^ revealed sustained rhythmic brain responses after sequence offset, produced only by intelligible speech, in the sensor cluster marked with plus signs. The “entrainment echo” was quantified as rate-specific responses (RSR; see Methods). T-statistics were obtained from a t-test against 0 (reflecting the null hypothesis). **C.** RSR in the cerebellum as a function of time, for the cluster shown in D. Time 0 corresponds to the first omitted word in the sequence. The horizontal lines show time points with significant RSR (FDR-corrected). The shaded areas correspond to time-windows of interest to test for entrained (brown) and sustained (blue) rhythmic brain responses, defined in the original study^9^. **D.** Source-localised t-statistics (RSR against 0) for sustained rhythmic responses in the intelligible condition (the equivalent on the sensor level is shown in B). Voxels included in the statistically significant cluster in the cerebellum are shown in brighter colours, the other voxels are faded out. **E.** As in D, but for the contrast between intelligible and unintelligible speech. Panels A and B are reproduced from the original publication^9^.

The original study^9^ did reveal rhythmic entrainment echoes in the MEG that were strongest at posterior sensors (Fig. 1B). Source-level analyses, however, were restricted to the cortical surface and did not produce clear results. Here, we hypothesized that the cerebellum is a driving force behind the observed sensor-level echoes and involved in the update of temporal prediction that is required when a stimulus stops^3^.

### Neural entrainment in the cerebellum and its echo

Based on the original study^9^, we computed an index that quantified rhythmic brain responses specific to the stimulus rate. This measure was labelled rate-specific response (RSR) and was obtained by contrasting rhythmic responses (quantified using inter-trial coherence, ITC) at a frequency that corresponds to the stimulus rate, with those at the same frequency but during stimulation at a different rate (see Methods). An RSR that is reliably larger than 0 during or after the rhythmic sounds reflects an entrained response or its echo, respectively (see Methods and ref^9^). We computed the RSR on the source level with a focus on the cerebellum.

We first tested whether activity in the cerebellum entrains to the stimulus rhythm. The original study, along with previous work^9,16–18^, already demonstrated entrained activity in auditory, motor, and inferior frontal cortical areas. Here, we found that this network includes the cerebellum. During presentation of rhythmic sounds, RSR was reliably larger than 0 in a cluster that comprised the whole cerebellum (Fig. S1). This was the case for intelligible (p = 0.0003, summed t = 136410, 22375 voxels in cluster) and unintelligible speech (p = 0.0003, summed t = 141530, 22375 voxels in cluster). Entrainment (i.e., RSR > 0) remained significant and constant in the cerebellum for all time points preceding the first omitted word (time 0 in Fig. 1C; horizontal lines correspond to FDR-corrected p < 0.05). However, it did not differ between intelligible and unintelligible speech (no significant cluster). This is in contrast to entrainment in auditory and inferior frontal areas, where an advantage for intelligible speech was described before, including in the original study^9,16,19^. Together, the cerebellum entrains to rhythmic speech sounds irrespective of their intelligibility.

We next tested whether entrained activity in the cerebellum persists after a sequence offset – the hypothesized “entrainment echo”. Fig. 1D shows the source-localized RSR (as t-statistic; see Methods), averaged within the silent period after intelligible speech. We found a cluster of voxels in the cerebellum with a reliable RSR after intelligible (p = 0.048, summed t = 9417.4, 3989 voxels in cluster) but not unintelligible speech (no significant cluster). This cluster comprised areas Crus II and Crus I and showed a reliably stronger RSR after intelligible than after unintelligible speech (Fig. 1E; p = 0.039, summed t = 11446, 4579 voxels). Numerically, the entrainment echo in the Crus II/I cluster was strongest throughout the brain (Figs. 1D,Eshow RSR values for all voxels, including those not included in the statistical analysis and therefore faded out). The time-resolved RSR (Fig. C) further demonstrates that this echo was not driven by the omission of an expected stimulus (time 0 in Fig. 1C) but concerned the whole silent period between stimulus sequences. Note again that the speech-specific entrainment echo, visible in Fig. 1C, is not preceded by speech-specific entrainment in the cerebellum. Together, these results suggest that the sensor-level echo (Figs. 1B) found previously^9^ is driven by the cerebellum and only present after intelligible speech.

### Sustained rhythmic responses in the cerebellum drive left inferior frontal gyrus (IFG)

We then investigated whether rhythmic responses in the cerebellum interacted with activity in other brain regions. We focused on the cerebellar region with the strongest sustained response (bilateral Crus II), and tested its directed connectivity (see Methods) with two other regions of interests (ROI), selected based on their strong entrained response in the same dataset (Fig. 2C in ref^9^). These ROI were bilateral superior temporal gyrus (STG) and inferior frontal gyrus (IFG, pars opercularis), both known to contribute to speech perception^20–22^. Responses were averaged across left and right Crus II due to their proximity and highly correlated activity (e.g., r = 0.76, p = 0.0001 for RSR in intelligible speech). As an equivalent to RSR, we computed a measure of connectivity that is only different from 0 if the underlying connectivity is specific to the stimulus rate (rate-specific connectivity, RSC; see Methods). In this case, the sign of RSC reflects the directionality (ROI A driving ROI Bor vice versa).

**Figure 2.**
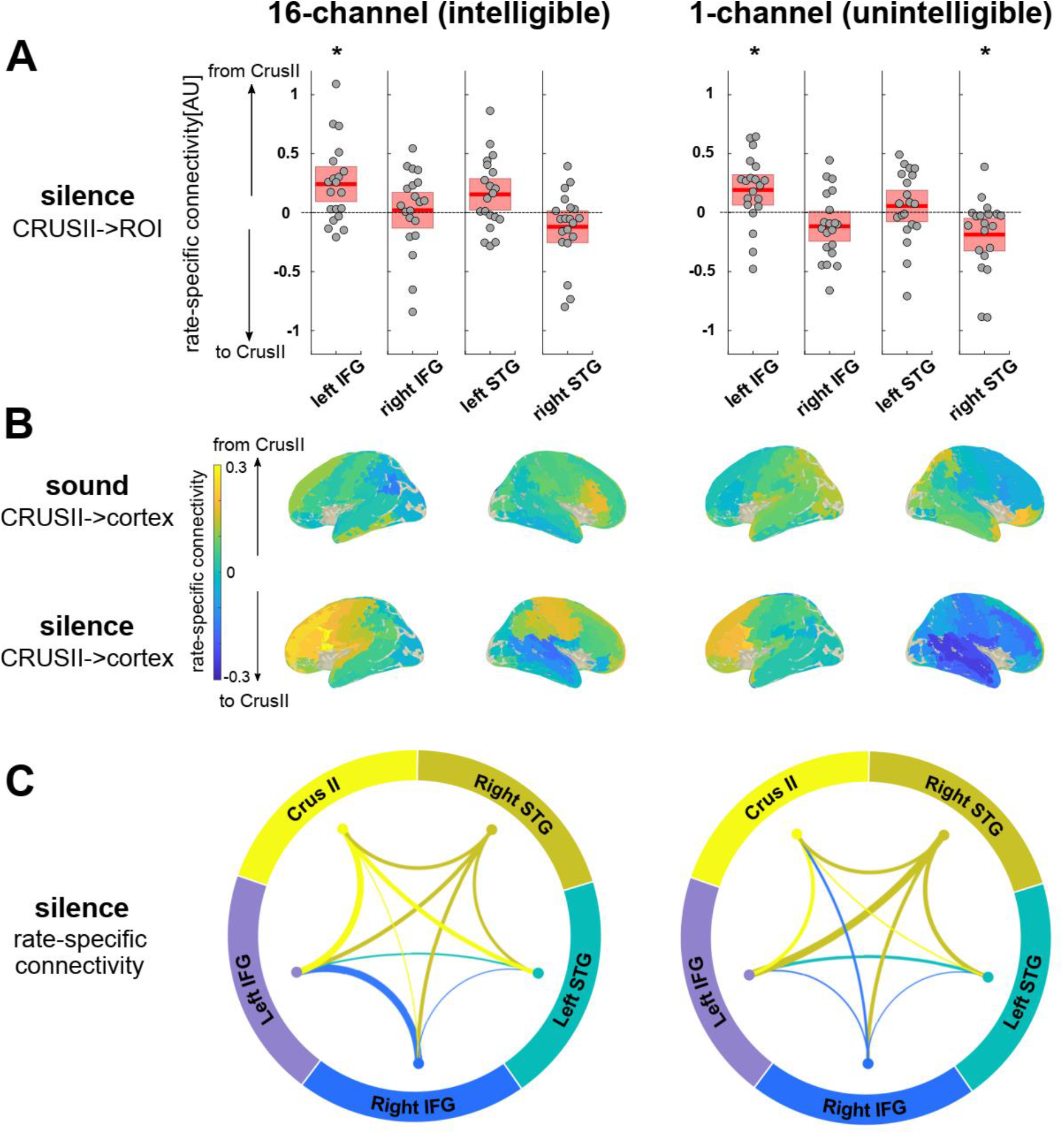
Rate-specific connectivity (RSC) results, for intelligible (left column) and unintelligible speech (right column). **A.** RSC during the silent period (i.e., during the entrainment echo) for connections between Crus II and two bilateral cortical ROIs. Points represent data from individual participants. Red lines and areas show mean and standard deviation, respectively. The asterisks indicate an RSC reliably different from 0 (dashed line). B. RSC between Crus II and other cortical areas during rhythmic sounds (top row) and silent period (bottom row). The average RSC across participants is colour-coded. C. RSC between ROIs in the silent period. The line thickness indicates the strength of connectivity, the colour shows the dominant direction (one colour originates from each ROI).

Fig. 2A shows average RSC values and their distribution in the silent period, for connections between Crus II and each hemisphere of the other two ROIs. We found that Crus II reliably drives left IFG after both intelligible and unintelligible speech (intelligible: t(19) = 3.20, p = 0.005; unintelligible: t(19) = 2.94, p = 0.009; both FDR-corrected p < 0.05). In addition, we found that right STG drives Crus II, but only after unintelligible speech (intelligible: t(19) = -1.75, p = 0.10; unintelligible: t(19) = -2.65, p = 0.016; unintelligible FDR-corrected p < 0.05). In contrast, we did not find any reliable rate-specific connectivity during rhythmic sound presentation (all FDR-corrected p’s > 0.40). When contrasting RSC values across ROIs, conditions (intelligible vs unintelligible), and time periods (sound vs silence), we found a significant interaction between ROI and time period (repeated-measures ANOVA: F(3,57) = 7.5, p = 0.0004; significant main effect of ROI was not interpreted due to the significant interaction). This was driven by an RSC between Crus II and left IFG that was higher during silence than during sound, and an RSC between Crus II and right IFG as well as between Crus II and right STG that was higher during sound than during silence.

Our principal hypothesis focused on the cerebellum and its connectivity with two cortical ROIs involved in speech processing. In less constraint analyses, Figs. 2B,C explore and illustrate the reported effects further. Fig. 2B shows RSC between Crus II and other cortical regions during (top row) and after (bottom row) intelligible (left column) and unintelligible speech (right column). During the entrainment echo, the left IFG was indeed the cortical region that was driven most strongly by Crus II. Acluster-based permutation test revealed a cluster for intelligible speech that comprised left IFG. However, it did not reach conventional statistical significance due to the larger number of areas included (p = 0.10, summed t = 8.38, 3 connections in cluster; no positive cluster for unintelligible speech). The right hemisphere was dominated by negative RSC values (reflecting connectivity with Crus II), in particular after unintelligible speech and in a superior temporal cluster (p = 0.13, summed t = -7.00, 3 connections in cluster).

Fig. 2C gives a more complete picture of the connectivity patterns between ROIs during the entrainment echo. After intelligible speech (left panel), apart from connectivity between Crus II and left IFG described above, the right IFG drove the left IFG (t(19) = 3.35, p = 0.003; FDR-corrected p < 0.05). In contrast, this bilateral connection was not present after unintelligible speech (right panel). Instead, the left IFG was driven by the right STG (t(19) = 3.57, p = 0.002; FDR-corrected p < 0.05).

In sum, we found rate-specific connectivity patterns during the entrainment echo, dominated by Crus II driving activity in left IFG. This result was common to both intelligible and unintelligible speech whereas the latter involved additional recruitment of the right STG.

## Discussion

When a rhythmic stimulus stops, entrained neural activity persists for some time. This effect, termed “entrainment echo” or “forward entrainment”^9,11,23–26^, is a hallmark of endogenous oscillations that have been entrained by a stimulus rhythm^15,27^. Entrainment echoes also provide interesting insights into neural processes that are activated when temporal predictions (induced by a rhythmic stimulus) are violated and need to be updated. Indeed, initial theories considered temporal prediction as one of the primary functions of neural entrainment^28,29^.

The current results revealed that rhythmic intelligible speech produces the strongest entrainment echoes in the cerebellum (Fig. 1D). During rhythmic speech, cerebellar activity entrained to the stimulus rhythm, but the entrainment was reduced compared to that in auditory or frontal cortical regions. This observation is in line with findings that the cerebellum is active when a stimulus rhythm *changes*, rather than during the rhythm itself^3,5^. Further evidence shows also that patients with cerebellar lesions struggle to adapt their tapping to a beat when the latter changes its rate^3,5^. Here, we did not specifically measure temporal prediction. However, the presence of entrainment echoes in the cerebellum, together with the cerebellum’s role in time perception and temporal prediction^3,30–33^, suggests that entrainment echoes might be a neural signature of temporal prediction (or its error). In future studies, these echoes might therefore become an important variable for both the diagnosis and treatment of corresponding deficits, as evident in schizophrenia^34^, dyslexia^35^, autism^36^ or attention-related disorders^37^. Entrainment echoes in auditory perception only seem to occur after stimulation between 2 and 8 Hz^11,26^. Another open question is whether similar rate-specificity can be found for cerebellar entrainment echoes, and whether these go along with rate-dependent changes in temporal predictions.

We also found that the cerebellar entrainment echo is not a local phenomenon but entails information exchange with cortical regions. This conclusion is based on rate-specific connectivity patterns (Fig. 2) that provided additional clues into the echo’s potential role in temporal prediction. Notably, we found that the cerebellum *drives* rhythmic activity in left IFG, rather than being driven. This finding suggests that the cerebellum provides the cortex with updates about expected event timing; in particular, it might signal errors between predicted and actual timing (*prediction error*^38^). This notion is in line with a recent study suggesting that the adaptation to dynamic environmental changes rely on “internal models” in the cerebellum and its connectivity with cortical areas^39^. It is also supported by a reported association between cerebellum and auditory hallucination^40^ (often considered to reflect false relay of prediction error^41^). Another possibility is that the echo in the cerebellum reflects a memory trace of the preceding rhythm that is necessary when it has to be compared with a new one. Such a process could be important to detect changes in the temporal structure of a stimulus. Speech perception is highly context-dependent, including the temporal domain: For instance, the perception of a word can depend on the rate of preceding speech^42^. Accordingly, a memory of preceding speech rates, in form of the entrainment echo, might also be necessary for successful speech perception. These hypothetical functional roles of entrainment echoes (relay of prediction error or memory trace) can also explain why rate-specific connectivity was only found after, but not during the presentation of rhythmic sounds: When prediction errors are low, or memory traces might not be needed when a stimulus does not change. Only when the stimulus rhythm stops or changes, the corresponding mechanisms might be activated and lead to the upregulation of connectivity that we observed.

A previous review^8^ has highlighted the lack of studies that examine the cerebellum’s role in speech processing by directly contrasting speech and non-speech stimuli. This was possible in the current work that compared responses to highly intelligible (16-channel) and unintelligible, noise-like (1-channel) vocoded speech. The use of noise-vocoding ensured very similar broadband amplitude envelopes^43^ for both stimulus types that might otherwise have biased results. In addition, entrainment echoes were not “contaminated” by motor activity, a common issue in studies targeting the cerebellum^8^.

By contrasting responses to intelligible and unintelligible speech, we revealed several interesting effects in the entrainment echo and corresponding cerebellar-cortical interactions. Firstly, entrainment echoes were stronger after intelligible speech and occurred in cerebellar subregions (Crus I/II) that have been associated with speech processing^8,44^. In contrast, echoes were not detectable after unintelligible stimuli (Fig. 1C). Although intelligible speech produces stronger neural entrainment in several cortical regions^9,16,17,19^, this was not the case in the cerebellum (Fig. 1C). It is therefore unlikely that the speech-specificity in entrainment echoes is merely due to a stronger sound-evoked response that persists after a stimulus offset, rather than a “true” advantage for intelligible speech that is specific to the echo. It also implies that entrainment echoes are not a simple continuation of a neural process that was already active during the rhythmic stimulus presentation. Secondly, the connectivity between Crus II and left IFG was stronger after intelligible speech, whereas the additional connection from right STG to left IFG, visible for unintelligible speech, was absent in this case (Fig. 2C). There are several possible explanations for this finding. Based on its acoustics, 1-channel vocoded-speech can be considered more rhythmic than 16-channel speech as all spectral frequencies are co-modulated (i.e., modulated by the same amplitude envelope). It is possible that auditory regions (such as STG) are recruited more readily by such acoustic rhythms. Rhythmic irregularities in speech are easier to detect when it is intelligible^45^, possibly due to the presence of linguistic features that help predicting upcoming information. When speech becomes unintelligible, temporal predictions cannot capitalise on intelligibility anymore and might instead rely on acoustically driven information. This might explain why the primary “driver” of left IFG changed from Crus II to right STG for unintelligible speech (see ref^8^ for a similar idea). Finally, it has been hypothesized that the cerebellum is involved in predictions that are based on learnt associations, rather than new ones that involve the cortex^8^. As participants know and understand intelligible but not unintelligible speech, this notion can also explain the current results. Together, these findings support an important role of the cerebellum, in particular Crus II, in speech processing, and the existence of a mechanism that is tailored to intelligible speech, rather than acoustic information in general.

Despite these differences in entrainment echoes and their connectivity, these results also revealed commonalities between intelligible and unintelligible speech. In particular, rate-specific connectivity analyses revealed left IFG as an information “receiver” in both cases. The IFG, in particular the left IFG, is an important hub for speech processing and strongly connected with auditory and motor regions^17,46^. It is also implicated in predicting upcoming speech^47^ and non-speech sounds^48^. A recent study^49^ concluded that the left IFG “supports prediction reconciliation in echoic memory” during speech perception. This conclusion is well in line with the current results and might explain why it received information from Crus II and STG when expectations were violated and such reconciliation is necessary. The echo from the cerebellum might then represent some kind of echoic memory, as already detailed above.

## Conclusion

Here we show that rhythmic intelligible speech produces rhythmic entrainment echoes in the cerebellum. These echoes might reflect updated temporal predictions that are relayed to left IFG and supported by right STG if the stimulus is not intelligible. These results support the idea that the cerebellum is involved in speech processing and temporal prediction in general.

## Methods

This study relies on a re-analysis of data acquired by van Bree et al.^9^. In the following, we summarise the experimental design and initial signal processing steps and refer to the original publication for further detail (Experiment 1 in that paper). We then provide a detailed description of analyses that are specific to the current study (source-level and connectivity analyses).

### Participants

We included data from 20 participants (10 females; mean ± SD, 37 ± 16 years) in the current analysis. The original study^9^ was approved by the Cambridge Psychology Research Ethics Committee (application number PRE.2015.132) and carried out in accordance with the Declaration of Helsinki.

### Stimuli and Experimental Design

Participants listened to (linguistically unrelated) monosyllabic words that were initially spoken to a metronome beat (inaudible to participants) and then combined into rhythmic sequences (Fig. 1A). These sequences were 2 or 3 seconds long and presented at one of two different rates (2 or 3 Hz). Speech sequences were noise-vocoded^43^ to manipulate their intelligibility, yielding intelligible 16-channel vocoded speech or unintelligible, noise-like 1-channel vocoded speech. Stimuli were presented through insert earphones that were connected to a pair of magnetically-shielded drivers.

Participants were asked to listen to the rhythmic sequences, and to press a button when they detected an irregularity in the rhythm of the sequence (red in Fig. 1A) that was present in 12.5 % of sequences. While they performed the task, participants’ MEG was recorded, combined with electroencephalography (EEG) that was not included in the present re-analysis. They completed a total of 640 trials. In each trial, intelligibility (16- or 1-channel speech) and duration (2 or 3 s) of the sequence was selected pseudo-randomly. Rate (2 Hz or 3 Hz) was kept constant within a block and selected pseudo-randomly for each of ten blocks. Each sequence was followed by an interval that was silent and used to measure sustained oscillatory responses (“entrainment echoes”). This interval was 2+x s long, where x corresponds to 1.5, 2, or 2.5 times the period of the sequence rate (i.e. 0.75, 1, or 1.25 s in 2-Hz blocks, and 0.5, 0.666, or 0.833 s in 3-Hz blocks). x was set to 2 in 50 % of the trials.

### MEG Data Acquisition and Pre-processing

A VectorView system (Elekta Neuromag) was used to collect MEG data in a magnetically and acoustically shielded room. This system has one magnetometer and two orthogonal planar gradiometers at each of 102 positions within a hemispheric array. Data were acquired at a sampling rate of 1 kHz and band-pass filtered between 0.03 and 333 Hz. Five head-position indicator (HPI) coils and two bipolar electrodes were used to monitor head position and electrooculography activity, respectively. A 3D digitizer (FASTRAK; Polhemus, Inc.) was used to record the positions of the HPI coils, and ∼70 additional points evenly distributed over the scalp relative to three anatomical fiducial points (the nasion and left and right preauricular points). The temporal extension of Signal Source Separation^50^ in the MaxFilter software (Elekta Neuromag) was applied to suppress sources of noise, compensate for motion, and reconstruct any bad sensors. MEG data were further processed using the FieldTrip software^51^ as well as custom-built scripts, both implemented in MATLAB (The MathWorks, Inc.).

### MEG Signal Processing and Statistical Analyses

Rhythmic MEG responses during and after the rhythmic sequences were quantified using intertrial phase coherence (ITC). ITC of a signal (at a given frequency and time) is high if its phase is consistent across trials^52,53^, and ranges between 0 (no phase consistency) and 1 (perfect phase consistency). Here, this phase was estimated in sliding time windows of 1 s (step size 20 ms), using Fast Fourier Transform (FFT). Example ITC values are visualised in Fig. 1E of the original publication^9^. From these time-resolved ITC values, we selected two time-windows of interest: One that captures the neural response *during* the rhythmic sounds, but avoids sequence on- and offsets (-1 s to -0.5 s relative to the first omitted word; shaded brown in Fig. 1C); the other immediately after the rhythmic sounds, but avoiding their offset (+0.5 s to +2 s relative to the first omitted word; shaded blue in Fig. 1C). In the original study^9^, entrainment echoes at the sensor level were found in this latter window and are reproduced in Fig. 1B.

ITC values were source-localised according to the following steps. Data from gradiometers and magnetometers were combined. (1) Co-registration of MEG data with individual T1-weighted structural MRI scans, based on realignment of the fiducial points. (2) Construction of individual lead fields using a single shell head model. (3) Spatial normalisation of brain volumes to a template MNI brain, and division into grid points of 5 mm resolution. (4) Estimation of spatial filter for each participant using a linear constrained minimum variance beamformer algorithm(LCMV^54^). This filter is designed to isolate distinct anatomical generators of the acquired MEG signals and might therefore work best when all regions are most and simultaneously active. Accordingly, we only used data acquired during acoustic stimulation to estimate the filter (+0.5 s to + 2 s relative to sequence onset). Note that this time window is difficult to compare with those used to quantify rhythmic MEG responses (described above), as these include a spectral analysis step (FFT) that “smears” responses in time. (5) Application of individual spatial filters to Fourier-transformed single-trial data, separately for each of the two time-windows of interest (during and after rhythmic speech; -1 s to -0.5 s and +0.5 s to +2 s relative to the first omitted word, respectively). (6) Computation of ITC, using the spatially filtered, Fourier-transformed single-trials. For each of the two stimulus rates (2 Hz and 3 Hz), this step yielded one ITC value per neural frequency of interest (2 Hz and 3 Hz) and time window, and for each of 2982 voxels inside the brain.

Source-level ITC values were combined to a rate-specific response (RSR) index. This index was obtained by contrasting ITC at a neural frequency that corresponds to the stimulation rate with ITC at the same neural frequency, but during another stimulation rate:

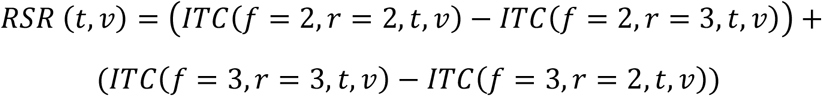

where *f* is the neural frequency for ITC, *r* is the sequence rate, and *t* and *v* are the time and voxel for which RSR is calculated, respectively. Note that an RSR larger than 0 reflects a rhythmic response that follows the stimulation rate at that time *t* and in voxel *v*. To test for reliable rhythmic neural responses, we therefore compared the RSR against 0, using Student’s t-test (one-tailed, reflecting the one-directional hypothesis). Given our study aim, we restricted this analysis to voxels in the cerebellum. We used an atlas-based parcellation for this purpose (AAL-Atlas^55^ in FieldTrip). We tested for rate-specific rhythmic responses separately for the two time-windows of interest and averaged RSR within each window. An RSR that reliably exceeds 0 in the first time window (during rhythmic sounds) demonstrates cerebellar activity aligned to the stimulus rhythm. An RSR that reliably exceeds 0 in the second time window (after rhythmic sounds) demonstrates that this rhythmic activity outlasts the stimulus at a frequency that corresponds to the stimulus rate, an effect that was termed “entrainment echo” and a hallmark of entrained endogenous oscillations^15,27^.

Statistically reliable effects were then determined by means of cluster-based permutation tests^56^ (5,000 permutations). Clusters (Fig. 1D,E) were considered significant if the probability of obtaining their cluster statistic (sum of t-values) in the permuted dataset was < = 5%. Corresponding effects in time (Fig. 1C) were corrected for multiple comparisons using False Discovery Rate (FDR).

To test for connectivity between the cerebellum and other brain regions, we first selected the part of the cerebellum with the strongest entrainment echoes in the analysis (bilateral Crus II; see Results). We then selected two bilateral regions of interest (ROI) that responded strongly to the rhythmic sounds in the initial study (Fig. 2C in ref^9^) and are known to be involved in speech processing^20–22^: the Superior Temporal Gyrus and Inferior Frontal Gyrus (pars opercularis).

We used spatial filters to construct time series for each dipole orientation in source space, resulting in three time series per voxel. Atlas-based parcellation was then applied to reduce the data to the 6 (3 bilateral) ROIs. Using singular value decomposition (SVD), the three strongest components (explaining most of the variance) from all dipoles in each ROI were extracted. From the resulting time series, we again selected two time-windows of interest, one centred on the rhythmic speech sequences (-1 to 0 s relative to the first omitted word), the other on the subsequent silent period (+0.5 to +1.5 s relative to the first omitted word). Note that these time-windows were selected to be one second long so that the output from the subsequent FFT includes estimates at 2 and 3 Hz, the two frequencies that were contrasted to quantify rate-specific responses and connectivity. The selected data were then subjected to a directional FFT-based connectivity analysis, to obtain the cross-spectral density (CSD) matrix and subsequent computation of multivariate nonparametric Granger causality^57^. We used multivariate Granger causality (mGC) measures to compute the directed influence asymmetry index (DAI) between two ROI:

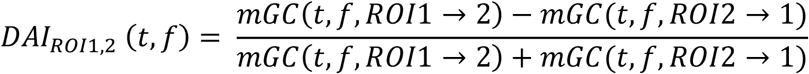

where *t* corresponds to one of two time-windows of interest (during or after rhythmic sounds; -1 s to 0 s and +0.5 s to +1.5 s relative to the first omitted word, respectively), *f* is the neural frequency (from the spectral transformation), and the arrows refer to the direction of mGC between the two ROI. The sign of DAI reflects the dominant direction of information flow between the two ROIs. Finally, we computed rate-specific connectivity (RSC) measures by following the rationale desribed for RSR above:

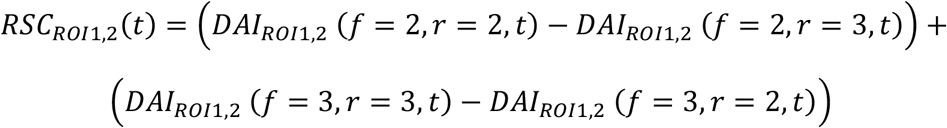

*r* is the sequence rate during (or after) which the DAI was observed. An RSC that is reliably different from 0 indicates connectivity between regions that occurs specifically at a frequency that corresponds to the stimulation rate. As for DAI, its sign reflects the dominant direction of information flow. For statistical analyses, we compared the RSC against 0, using Student’s t-test (two-tailed, reflecting the two-directional hypothesis). Prior to this test, RSC was averaged across left and right Crus II. This was done as they are difficult to distinguish in the MEG, due to their close proximity and evidenced by a high correlation between their values (e.g., r = 0.76, p = 0.0001 for RSR during intelligible speech). FDR was used to correct for multiple comparisons.

## Acknowledgements

Benedikt Zoefel is supported by grants from the Agence Nationale de la Recherche (ANR-21-CE37-0002) and Fondation pour l’Audition (FPA-RD-2021-10). Joachim Gross is supported by grants from the DFG (GR 2024/11-1, GR 2024/12-1, GR 2024/5-1). Sonja Kotz is supported by a grant from the Bial Foundation (102/22). The authors thank Sander Van Bree, Ediz Sohoglu, and Matthew H Davis for their contribution to the original study. This original work was supported by the European Union’s Horizon 2020 research and innovation programme under the Marie Sklodowska-Curie grant agreement number 743482 (Benedikt Zoefel), the British Academy/Leverhulme Trust (grant number SRG18R1\180733) (Benedikt Zoefel), and the Medical Research Council UK (grant number SUAG/044 G101400) (Matthew H Davis).

## Supplementary Figures

**Figure S1.**
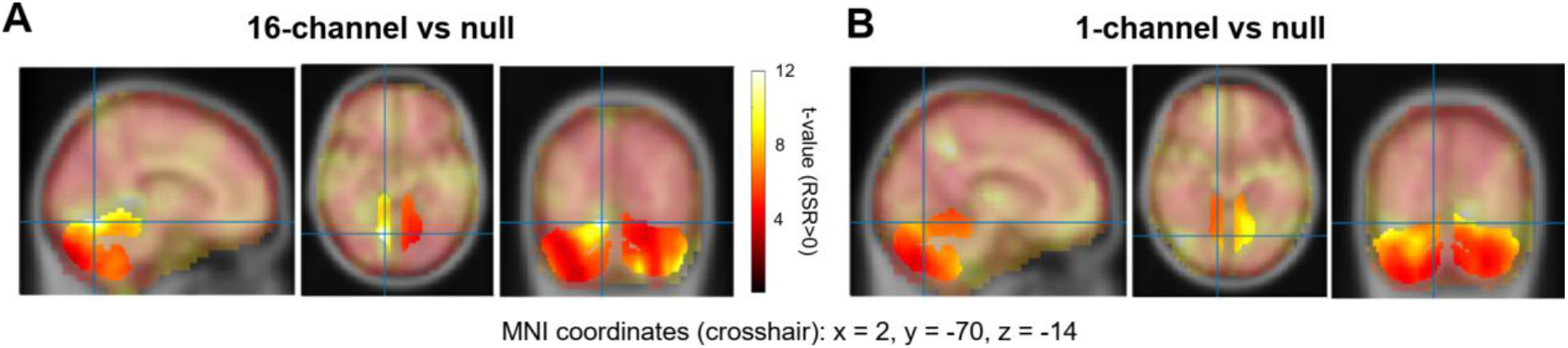
Source-localised RSR (t-statistics) during intelligible (A) and unintelligible (B) speech. An RSR > 0 reflects an entrained brain response. Voxels in the cerebellum are shown in brighter colours. Given our hypothesis, only the cerebellum was included in the statistical analysis (t-values are shown for other voxels only for completeness), revealing a significant cluster in the whole cerebellum.

